# Revisiting Hox gene evolution and Hox cluster linkage across Nematoda

**DOI:** 10.1101/2023.10.16.562615

**Authors:** Joseph Kirangwa, Dominik R Laetsch, Erna King, Lewis Stevens, Mark Blaxter, Oleksandr Holovachov, Philipp Schiffer

## Abstract

Hox genes are central to metazoan body plan formation, patterning and evolution, playing a critical role in cell fate decisions early in embryonic development in invertebrates and vertebrates. While the archetypical Hox gene cluster consists of members of nine ortholog groups (HOX1-HOX9), arrayed in close linkage in the order in which they have their anterior-posterior patterning effects, nematode Hox gene sets do not fit this model. The *Caenorhabditis elegans* Hox gene set is not clustered and contains only six Hox genes from four of the ancestral groups. The pattern observed in *C. elegans* is not typical of the phylum, and variation in orthologue set presence and absence and in genomic organisation has been reported. Recent advances in genome sequencing have resulted in the availability of many novel genome assemblies in Nematoda, especially from taxonomic groups that had not been analysed previously. Here, we explored Hox gene complements in high-quality genomes of 80 species from all major clades of Nematoda to understand the evolution of this key set of body pattern genes and especially to probe the origins of the “dispersed” cluster observed in *C. elegans*. We also included the recently available high-quality genomes of some Nematomorpha as an outgroup. We find that nematodes can have Hox genes from up to six orthology groups. While nematode Hox “clusters” are often interrupted by unrelated genes we identify species in which the cluster is intact and not dispersed.

## Introduction

Hox genes are crucial regulators of body patterning and development in bilaterian animals, controlling the identity and position of body structures along the anterior-posterior axis^1^. These body patterning genes were initially discovered in *Drosophila melanogaster* fruit flies through the striking phenotypes they gave rise to when genes were mutated^2^. In these homeotic phenotypes, one part of the body along the anterior-posterior axis is transformed into the likeness of another part of the body, such as Antennapedia mutations that transform fly antennae into legs^3^. In many bilaterian animals, these genes are organised into clusters in the genome, with the order of the genes reflecting their spatial expression during embryonic development (called collinearity)^2^. The last common ancestor of protostomes and deuterostomes is thought to have had a cluster of at least seven Hox genes characterised by a common transcriptional orientation and by colinearity in the order of the genes and their expression domains along the anterior-posterior axis^4–6^.

Vertebrate Hox gene clusters have a distinctive structural and transcriptional organisation. This is true of all of the Hox cluster copies present following the whole genome duplications in vertebrates^7,8^. Despite the generally larger size of vertebrate genomes, vertebrate Hox genes are very densely packed and Hox cluster span is much smaller than in many invertebrates. The clusters have low repeat densities, and while protein-coding genes unrelated to the Hox family are excluded, small RNA genes which regulate cluster expression can be interspersed. Indeed, clustering is strongly associated with the mechanisms of cross- and co-regulation of Hox gene expression^9,10^.

Clustering and collinearity are not always conserved. Successful animal taxa have managed to escape the constraint of having a tightly regulated, compact and complete Hox cluster. In several bilaterian phyla, Hox genes have undergone significant evolutionary changes including gene duplication, loss, and gain as well as changes in cluster organisation and expression patterns^11^. Some taxa have Hox clusters that are less tightly organised but still intact when compared to vertebrate counterparts, including the cephalochordates (Branchiostoma)^12^, sea urchins^13^, and insects (Apis^14–16^, Anopheles^17,18^ and *Tribolium castaneum*^19,20^). The Hox cluster in *Drosophila melanogaster* is split into two subclusters^21^. Splitting has occurred repeatedly and independently in other Drosophilids, at different positions along the cluster^16,22^. Disaggregation of the Hox cluster can be observed in tunicate chordates *Ciona intestinalis* (solitary tunicate) and *Oikopleura dioica* (appendicularian) where Hox loci are present but not tightly linked in the genome^23,24^.

By comparing the complements and developmental roles of Hox genes in different groups, it is possible to propose models of how the evolution of these genes has influenced the morphological diversity within various metazoan phyla. Loss of Hox genes can lead to body plan simplification while gene duplication can result in the acquisition of novel developmental processes^25,26^. Indeed the body plan of an animal can be a consequence of Hox gene loss as thought for the tardigrade *Hypsibius exemplaris*^26,27^. Hox gene loss is a common feature in Tardigrada, where a reduced complement of seven HOX genes including a local duplication of Hox9-13 was found^28^.

Nematomorphs are the closest relatives to Nematoda and are all parasitoids in arthropods. Both groups have very simple and vermiform body plan. A previous survey of Hox gene content using both PCR surveys and transcriptome analysis in *Paragordius* revealed a Hox gene complement of five ancestral ortholog groups. The presence of Hox2 (Pb) in *Paragordius* yet lost in the nematode counterparts revealed independent loss in Nematoda^28,29^. Recently published high-quality genomes from Nematomorpha (*Acutogordius australiensis, Nectonema munidae*, and *Gordionus montsenyensis*) show a relatively low number of genes (11,114, 8,717, 10,819 respectively) and lack a high proportion of universal single-copy metazoan orthologs^30,31^. Previous studies of Hox gene complement analysis in both Nematoda and Nematomorpha suffered from the lack of high-quality genomes. Therefore, a full complement of Hox gene evolution and their respective genomic organisation could not be well ascertained. These recently available high-quality genomes from both phyla provide a great resource to revisit the evolution of this key set of important developmental control genes.

Nematoda comprises ubiquitous and evolutionarily important organisms^32–35^ that have provided valuable insights into the genetics of development. The model *C. elegans* was considered a representative of the developmental biology and genomics of all Nematoda for many decades, but nematodes are highly diverse with over 28,000 described species^36,37^ and potentially more than 1 million species in the phylum^38,39^. They are found in a wide range of habitats including soil, freshwater and marine environments. Phylum Nematoda is split into three classes^32^: Dorylaimia (also known as Clade I), Enoplia (Clade II), and Chromadoria (Clade C). Within the largely marine Chromadoria, three speciose suborders within the largely terrestrial order Rhabditida are distinguished (Spirurina or Clade III, Tylenchina or Clade IV, and Rhabditina or Clade V). *C. elegans* is a member of Rhabditina (Clade V) in Figure 1. It has now become apparent that nematodes show substantial divergence in regard to cellular pattern formation in early development^40–42^, post-embryonic development^43–45^ and their general gene content^46–49^.

**Figure 1.**
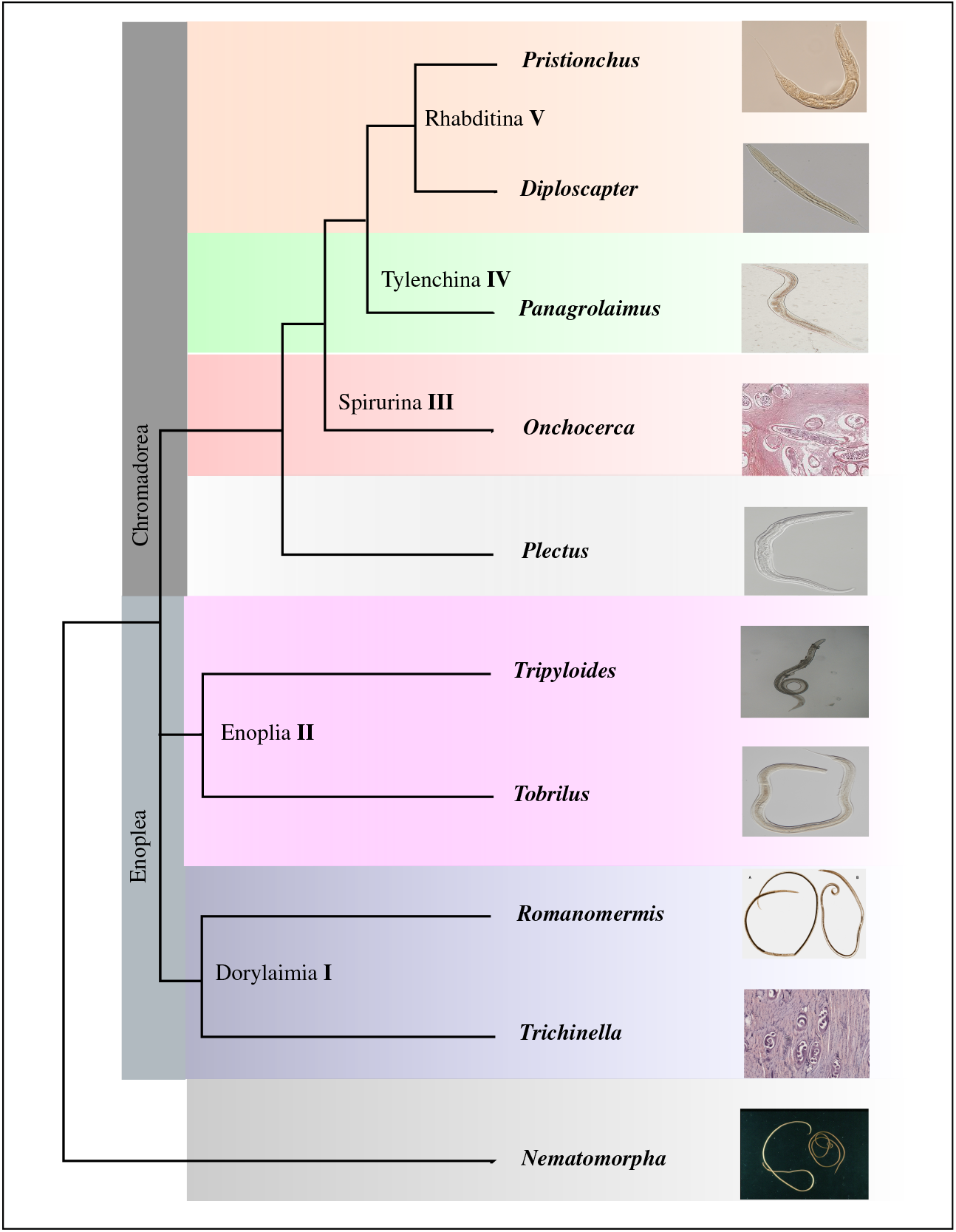
Cladogram of Phylum Nematoda showing Clades relevant to this work. The clade naming convention was introduced by *Blaxter et al.* in 1998.

The *C. elegans* Hox gene set has been shown to differ substantially from counterparts in other animal phyla. The six Hox genes include representatives of only four core Hox orthology groups (ceh-13 from the HOX1 group, lin-39 from HOX4, mab-5 from HOX6-8 and egl-5, php-3 and nob-1 from HOX9). The genes are linked but largely unclustered: they are distributed over an approximately >4 Mb span of *C. elegans* chromosome III, with many unrelated genes interspersed^25,50^. While there is some residual collinearity, the HOX1 and HOX4 orthologues of ceh-13 and lin-39 are inverted compared to the relative order of HOX6 and HOX9. In recent years it has become evident that many features of nematode embryonic development (including timings of cell division, cell lineage, and gene expression) are highly variable across the phylum^40,51^, and, like nematode genomes, evolve rapidly^29,48,52,53^. While Hox gene evolution has been well studied in *C. elegans* and a biased sampling of parasitic species from both Clade I and III^29^, our knowledge of Hox gene repertoire, Hox Cluster organisation and linkage in most nematode lineages is limited. Determination of the full complement of Hox genes in recently available nematode genomes across the Nematoda is fundamental to understanding the link between the genomic and morphological evolution of this phylum. Here we have analysed the Hox gene complements of a wide range of newly-sequenced, high-quality genomes from diverse nematodes to understand the evolution of this key set of body patterning genes.

## Results

### Hox gene complements in Nematoda

#### HOX gene evolution in nematodes is largely characterised by consecutive losses

Hox genes play critical roles early in embryonic development in determining cell fates so that the correct body parts are formed in the right places from nematodes to vertebrates. While it is possible to develop interim catalogues of Hox gene repertoires by directed PCR, transcriptomics or draft genome sequencing, to validate presence-absence patterns and explore synteny relationships high quality genome sequences are required. Across the analysed nematode species we identified loci attributable to six Hox orthology groups; HOX1 (arthropod lab, *C. elegans* ceh-13), HOX3 (zen), HOX4 (Dfd, lin-39), HOX6-8 (ftz, mab-5), HOX6-8 (Antp), HOX9-13 (AbdB, egl-5/php-3/nob-1). We did not find loci that were classifiable as HOX2 (pb), HOX5 (scr) and the Ubx/AbdA subtypes of HOX6-8 loci in any of the nematode genomes analysed. The maximum number of Hox loci in a single species was seven (in members of Spirurina / Clade III), and the minimum four (*Oncholaimus oxyuris*) (Figure 2). Loci attributed to HOX orthogroups HOX9-13 showed variable presence across the phylogeny.

**Figure 2.**
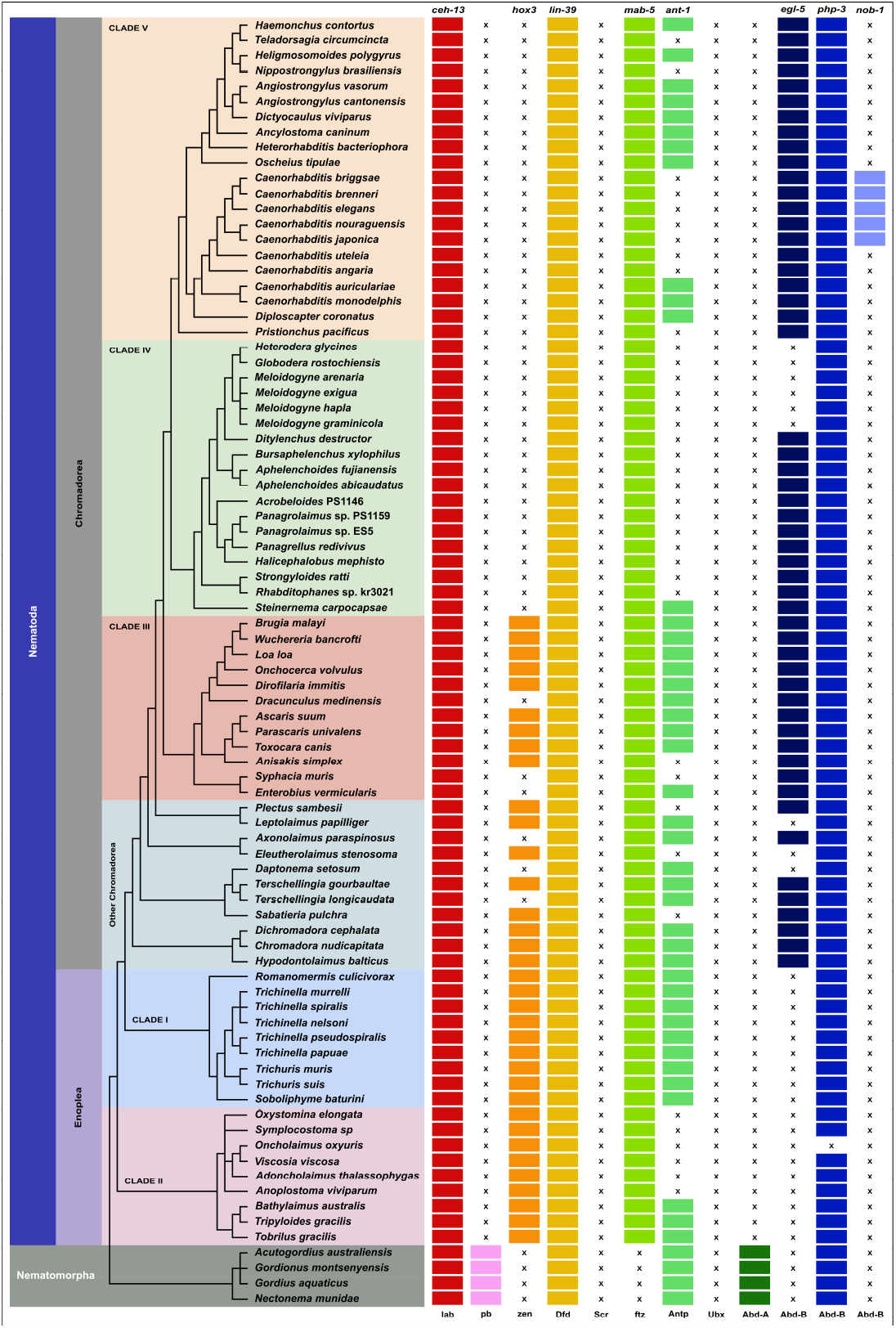
Nematoda and Nematomorpha Hox gene complements. Genes belonging to the same orthology group (HOX1-HOX9) are coloured similarly. X indicates the absence of the orthologue. The cladogram on the left shows the phylogenetic relationships of the nematodes analysed.

##### Our pipeline reliably recovers C. elegans HOX loci

The pipeline is dependent on a reference database consisting of highly curated HMM profiles for HOX homeodomains. These profiles are obtained through protein alignments from closely related species, where *C. elegans* and Amphioxus Hox homeodomains are employed as the in-group and outgroup, respectively. This choice holds particular significance due to the heightened sensitivity of sequence similarity search techniques to the reference database’s protein quality. Furthermore, the rapid evolutionary nature of nematode homeodomains adds to the rationale for this approach^29^. We applied the analysis pipeline (Figure 5) in order to test the ability to recover all the *C elegans* expected Hox loci. All the six expected loci were recovered (Figure 2).

##### Hox6-8 (Antp) loci variation in Rhabditomorpha (including Strongylomorpha=Strongyoidea which is an ingroup) crown group nematodes

Nematodes belonging to the infraorder Rhabditomorpha form a diverse group exhibiting various lifestyles, ranging from being free-living to parasitic organisms in animals. While *C. elegans* misses Hox6-8 (Antp) loci, we found presence in crown group nematodes except *Teladorsagia circumcincta* and *Nippostrongylus brasiliensis* which are parasites of sheep^54^ and rodents^55^ respectively. In the members of this group included in our analysis, all *C. elegans* Hox genes are present except for nob-1 (Figure 2).

##### nob-1 is a recent duplication within Caenorhabditis

Our study expanded to investigate the Hox gene complements in diverse *Caenorhabditis* nematodes beyond *C. elegans* (Figure 3). The Hox gene repertoire resembled that of *C. elegans*, with an exception: the detection of a recent nob-1 gene duplication^29^ in species belonging to the elegans and japonica subgroups including *C. briggsae, C. brenneri, C. nauraguensis*, and *C. japonica*. The nob-1 gene was absent in the earlier branching *Caenorhabditis* members of the Drosophilae subgroup (Figure 2). Among the three posterior Hox gene class (egl-5, php-3, and nob-1) in *C. elegans*, the same genes were present in close relatives. However, other *Caenorhabditis* species have two HOX9-13 loci corresponding to egl-5 and php-3, while no other nematode displayed more than two members of HOX9-13. The inclusion of *Diploscapter* as a very close outgroup provides strong support for the idea that nob-1 is unique to the *Caenorhabditis* genus. (Figure 2).

**Figure 3.**
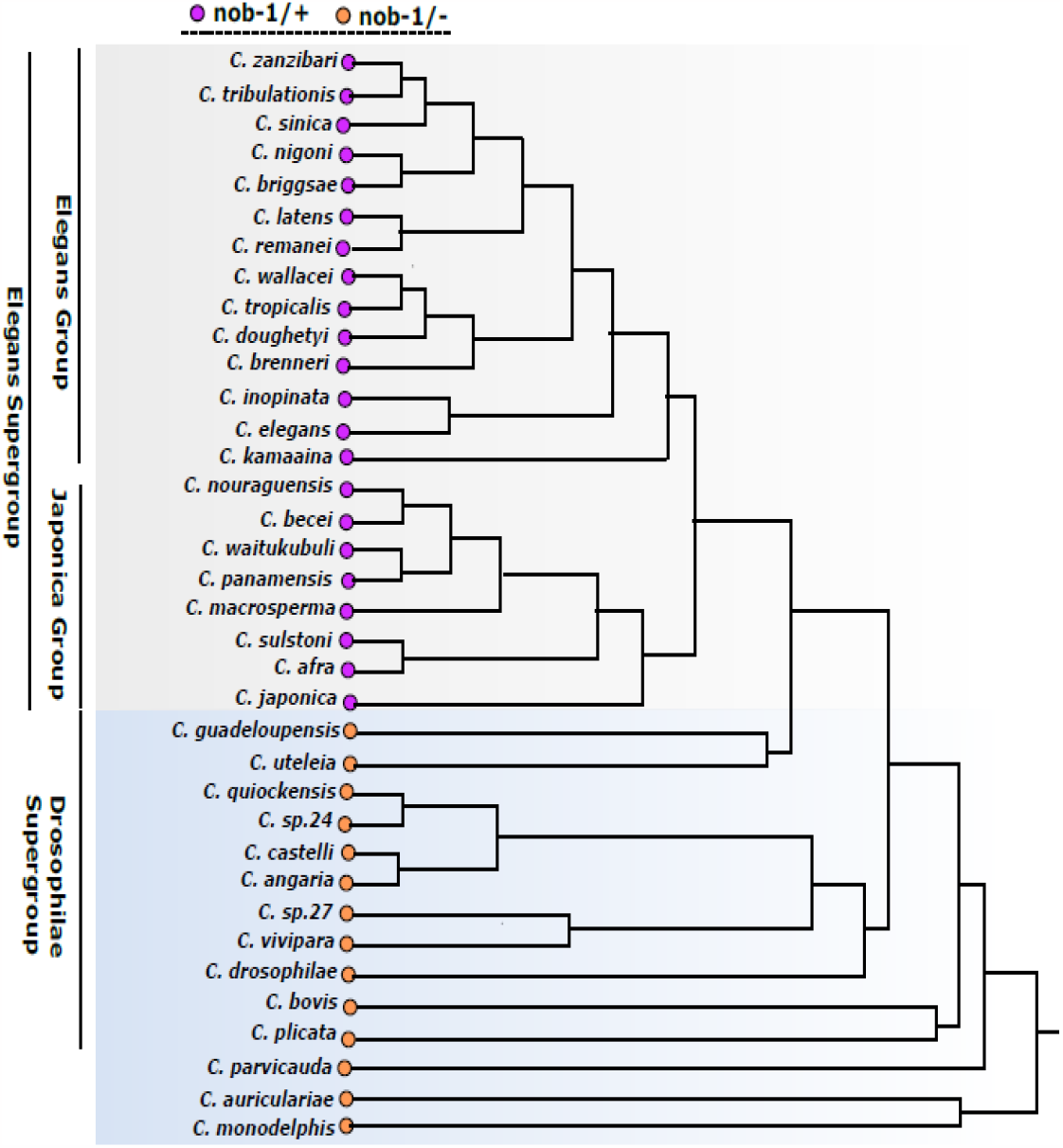
nob-1 Hox gene complement across *Caenorhabdtis* lineage.

##### Hox3 gene loss in Tylenchina and Spirurina

During our investigation, we noted that all the examined members within the Tylenchina clade lacked HOX3 loci. Within the Spirurina clade, specific instances of Hox3 loss were identified in *Syphacia muris, Enterobius vermicularis*, and *Dracunculus medinensis*, which are illustrated in Figure 2. A single loss of ant-1 at the base of Tylenchina, except in the case of *Steinernema carpocapsae*, as depicted in Figure 2. However, despite the presence of ant-1 loci in all Spirurina species included in our study, *Syphacia muris* and *Anisakis simplex* were exceptions where this locus was found to be absent. Furthermore, we observed separate instances of Hox9-13 (egl-5) loss among the members of the Tylenchoidea crown group.

##### Hox gene complements in early-branching chromadoreans

Previous investigations into the Hox gene complement among members of “Clade C” in Chromadorea were limited due to insufficient genomic resources, impeding a comprehensive understanding of Hox gene evolutionary patterns^28^. In this study, we expanded our analysis by utilizing newly available draft genomes in early-branching Chromadorea. In certain species, we observed Hox3 loss (*Terschellingia longicaudata, Daptonema setosum*, and *Axonolaimus paraspinosus*). Furthermore, the Antp-like locus was absent in *Plectus sambesii, Sabatieria pulchra*, and *Eleutherolaimus stenosoma*. Additionally, we identified independent cases of Hox9-13 (egl-5) loss in *Leptolaimus papilliger, Daptonema setosum*, and *Eleutherolaimus stenosoma*. Other members of this Clade collectively possessed a total of seven Hox genes, as depicted in Figure 2.

##### An independent loss of ant-1 locus in Enoplida, and a single HOX9-13 (php-3) in Enoplean nematodes

The Clades I and II (*Enoplea*) class encompasses the earliest-branching lineages within the Nematoda. Previous investigations into the Hox gene composition of Enoplea were limited by the lack of genomic data for several important enoplean lineages. In this study, we delve deeper into the analysis, utilizing 9 genomes from Enoplia (Clade II) and 9 genomes from Dorylaimia (or Clade I). Initially, our findings revealed the presence of a collective total of 6 Hox gene loci in most Enoplean members (depicted in Figure 2). Among these, a solitary HOX9-13 orthologue was identified, which was classified as php-3. However, *Oncholaimus oxyuris*, a species with only four Hox orthologue groups, lacked any HOX9-13 locus, specifically the one (php-3) that would be anticipated. It’s important to note that this assembly originated from a single specimen amplified for long-read sequencing, and the absence might stem from residual assembly issues rather than actual loss (as shown in Figure 2). The egl-5 locus was absent in both Enoplea and Dorylaimia species. Furthermore, we observed independent losses of the Antp-like locus within sampled Clade II (*Enoplida*) members among Enoplia (Figure 2).

##### Nematomorpha have six Hox loci

Nematomorpha, the sister phylum of Nematoda^56^, represent a group of parasitoid worms commonly known as horsehair worms or Gordian worms. Earlier investigations into the Hox gene complements within Nematomorpha yielded inconclusive results, primarily attributed to inadequate sequence data that hindered the assignment of numerous Hox genes to specific orthologue groups^29^. In our investigation of their Hox gene complements from 4 recently available genomes, we found Hox loci corresponding Hox1 (lab), Hox2 (Pb), Hox4 (Dfd), Hox6-8 (Antp), Hox6-8 (Abd-A) and Hox9-13 (Abd-B). We did not find Hox loci corresponding to Hox3 (zen) and Hox6-8 (Ftz) (Figure 2).

#### Hox cluster organization in the Nematoda

Clustering and collinearity are prominent features observed in the evolutionary path of HOX genes, and this pattern has proven to be consistent across a wide spectrum of species. In contrast to the loose clustering of Hox genes in three loci pairs on chromosome III in *C. elegans*, the analysis of nematode genomes in the present study reveals more condensed Hox gene clusters. These clusters exhibit variations in the number of genes, the sequence of gene arrangement, and the orientation of transcription. Furthermore, we observe interruptions within these Hox gene clusters by the presence of Non-Hox genes.

##### Unique characteristics of Hox gene organization in Bursaphelenchus

We conducted a comprehensive investigation into clustering, gene order, and orientation within Hox gene sets across the phylum. Our analysis revealed potential HOX clusters in several species. Notably, in the two *Bursaphelenchus* species, four out of the five Hox loci were found to be closely linked, with a separation of less than 40 kb in *B. xylophilus*, while the HOX9-13 locus, php-3, was separated by 300 or 700 kb. Within this core set of four loci, the genes maintained the same relative order, but in *B. okinawensis*, the core cluster exhibited an inversion with respect to the position of php-3 compared to its arrangement in *B. xylophilus*. The order of genes in the core did not align with the expected HOX1-HOX6-8 ordering (Figure 4).

**Figure 4.**
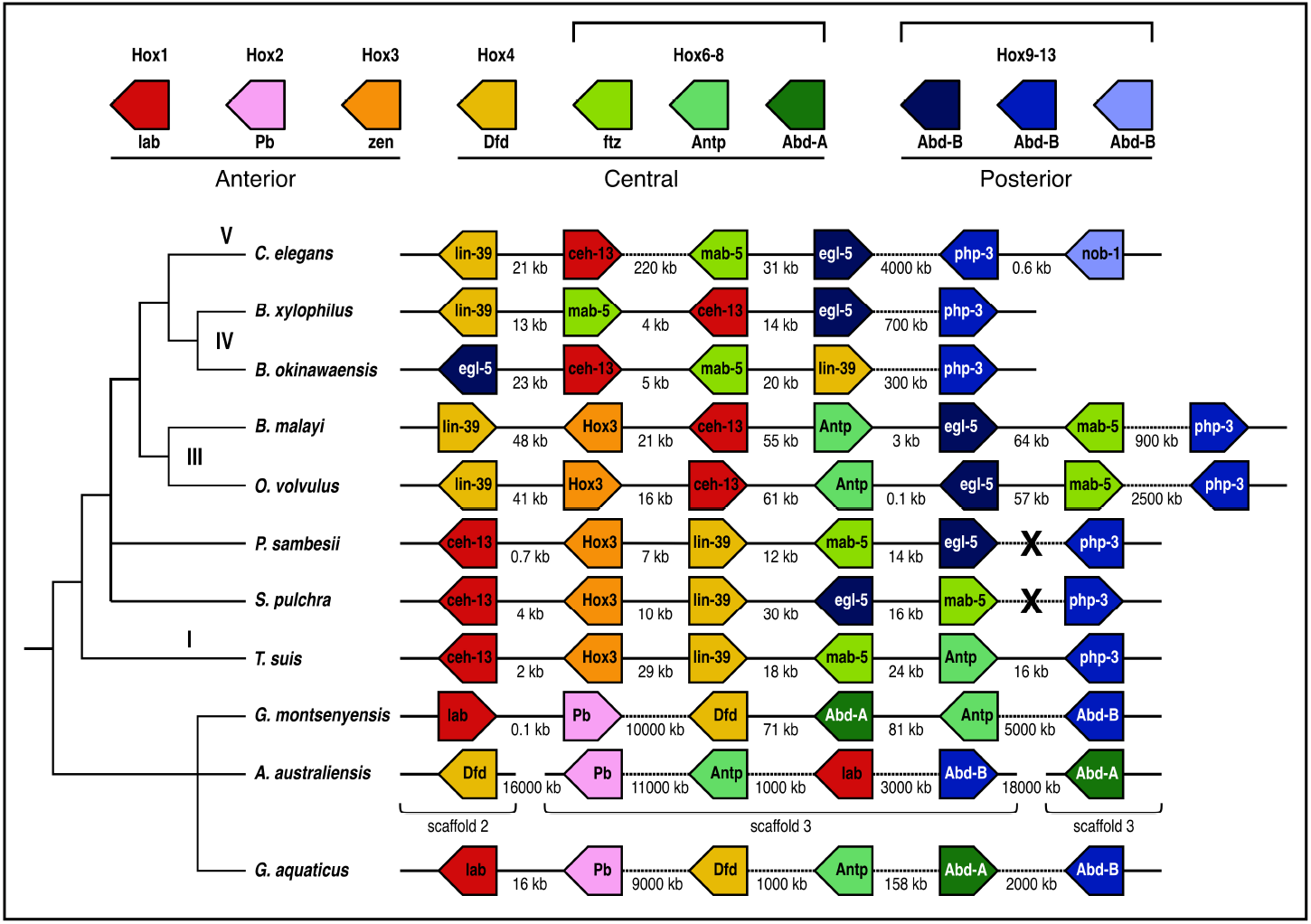
Nematoda and Nematomorpha Hox gene clusters. Synteny relationships of Hox cluster loci from eight nematodes and three nematomorphs are shown. Identified Hox genes and their respective transcriptional orientations are indicated by arrow boxes which are coloured by orthology group. Dashed horizontal lines indicate cluster breakage in scaffolded genomes. Lineage-specific duplications include paralog groups Hox6-8 and Hox9-13. Crosses indicate breaks that result from gaps in un-scaffolded draft genomes. The cladogram on the left summarises the relationships of the nematodes.

##### Some members of Spirurina display Hox clusters containing a greater complement of Hox genes

Both *Brugia malayi* and *Onchocerca volvulus* possess expanded Hox clusters containing seven linked Hox genes, of which six are closely clustered, and both species have a greater number of Hox genes compared to most other members of the Nematoda group. This core cluster comprises six members (ceh-13, hox-3, lin-39, mab-5, ant-1, egl-5), while the HOX9-13 php-3 locus is situated at a distance of 0.9 Mb (*B. malayi*) or 2.5 Mb (*O. volvulus*). Although the gene order within the core group remains consistent between the two species, it does not adhere to the typical HOX1-HOX6-8 structure, and there are differences in the transcriptional orientation of the Hox loci between these species (Figure 4).

##### Hox cluster organization in basal Chromadorea

In *Plectus sambesii* and *Sabatiera pulchra* (Chromadoria) clusters containing five Hox genes, the homologues of ceh-13, hox-3, lin-39, mab-5 and ant-1, were identified. The cluster in *P. sambesii* was particularly compressed (less than 40 kb) (Figure 4). The clusters differed by an inversion of the mab-5 and ant-1 pair, but in *P. sambesii* the order of the loci corresponds to the model structure, albeit the transcriptional order does not correspond (Figure 4). The HOX9-13 locus php-3 was located on a different genomic contig in both nematodes (Figure 4).

##### Hox loci in Trichuris are linked together in CiS and adhere to the model order

In the instance of *Trichuris suis*, a member of the *Dorylaimia* lineage, a cluster covering about 100 kb was detected, possessing six Hox loci that were interconnected and transcribed from distinct DNA strands (Figure 4).

##### In Nematomorpha, the arrangement of Hox clusters follows the common sequence of HOX1-HOX6-8 order

The Hox genes in *Gordionus montsenyensis* exist as a fragmented cluster (Figure 4); anterior paralogs (Hox1(lab) and Hox2(Pb)) are seperated from the three central group paralogs (Hox4(Dfd), Hox6-8(Antp) and Hox6-8(Abd-A)) by approximately 10 Mb (Figure 4). The posterior Hox9-13 (Abd-B) is further way on the same scaffold by more than 4 Mb (Figure 4). The direction of transcription of the loci is not ordered. The order of the genes in the core does reflects the expected HOX1-HOX6-8 ordering except that HOX6-8/AbdA and Hox6-8/Antp genes are inverted with respect to the order found in ancestral cluster.

*Acutogordius australiensis* has members of Hox genes present in 3 seperate genomic scaffolds. Scaffold_2 contains Hox4 (Dfd), scaffold_3 contains 4 Hox loci, scaffold_4 contains Hox6-8 (Abd-A). The order of the genes does not reflect the expected HOX1-HOX6-8 ordering. The direction of transcription of the loci is not ordered (Figure 4). Hox genes in *Gordius aquaticus* are linked on scaffold 5. The order of Hox loci reflects the expected HOX1-HOX6-8 ordering. The direction of transcription of the loci is not ordered (Figure 4).

#### Hox clusters in nematodes are interrupted by non-Hox genes

Although vertebrate genomes are typically larger, the arrangement of vertebrate Hox genes is notably compact, with Hox clusters spanning a smaller region compared to many invertebrates. These clusters exhibit low repeat densities and exclude protein-coding genes unrelated to the Hox family. One of the few gene types that are found inside the clusters is small RNA genes (microRNAs) that have roles in Hox gene regulation. This suggests a strong constraint on the Hox cluster in vertebrates, likely because of the intimate coregulation of the Hox loci. We explored whether similar constraint acts on nematode Hox clusters. We identified non-Hox genes between Hox loci in nematode species analysed (Table 1). Where Hox loci were tightly clustered, the number of genes was reduced, but even in the smallest clusters (*B. xylophilus, P. sambesiiT. muris*), non-Hox loci were common (Table 1).

**Table 1.**
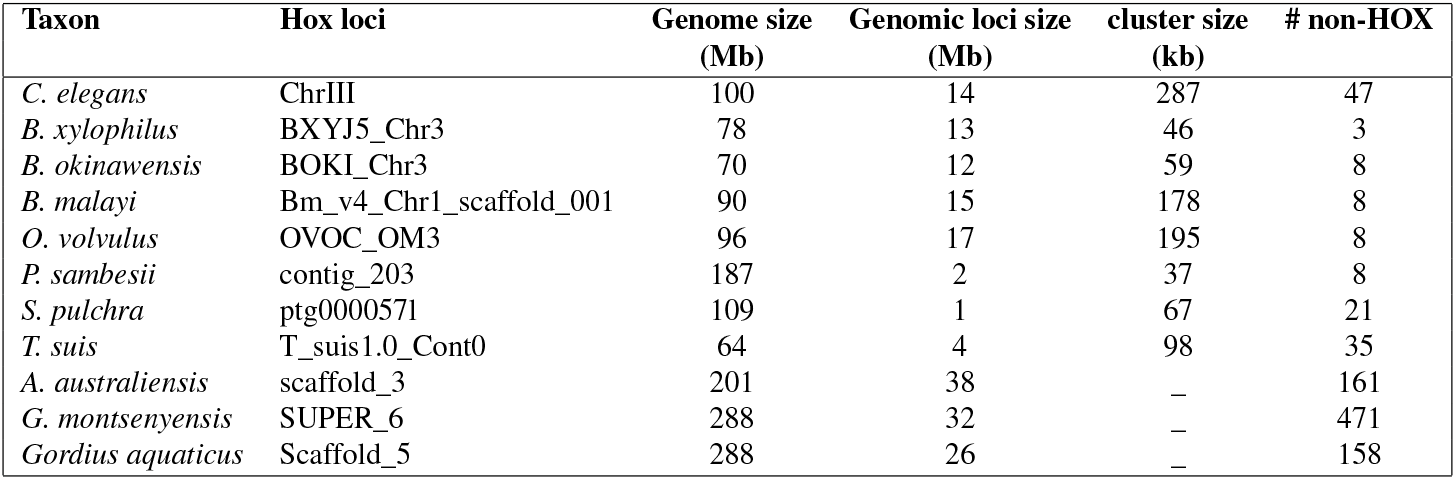
Hox cluster features of Nematoda and Nematomorpha from available high quality genomes.

| Taxon | Hox loci | Genome size<br>(Mb) | Genomic loci size<br>(Mb) | cluster size<br>(kb) | # non-HOX |
| --- | --- | --- | --- | --- | --- |
| *C. elegans* | ChrIII | 100 | 14 | 287 | 47 |
| *B. xylophilus* | BXYJ5_Chr3 | 78 | 13 | 46 | 3 |
| *B. okinawensis* | BOKI_Chr3 | 70 | 12 | 59 | 8 |
| *B. malayi* | Bm_v4_Chr1_scaffold_001 | 90 | 15 | 178 | 8 |
| *O. volvulus* | OVOC_OM3 | 96 | 17 | 195 | 8 |
| *P. sambesii* | contig_203 | 187 | 2 | 37 | 8 |
| *S. pulchra* | ptg0000571 | 109 | 1 | 67 | 21 |
| *T. suis* | T_suis1.0_Cont0 | 64 | 4 | 98 | 35 |
| *A. australiensis* | scaffold_3 | 201 | 38 | — | 161 |
| *G. montsenyensis* | SUPER_6 | 288 | 32 | — | 471 |
| *Gordius aquaticus* | Scaffold_5 | 288 | 26 | — | 158 |

## Discussion

Hox genes are key players in the structuring of animal body plans during development. The developmental role of Hox genes has been well studied in *C. elegans* and the second model nematode *Pristionchus pacificus*^25,29,57,58^. Our knowledge of the Hox gene repertoire, Hox Cluster organisation, and linkage in most nematode lineages is limited, with data restricted to some parasitic species scattered across Nematoda^29^. Most published nematode genomic resources come from Clades I, III, IV and V often missing a higher representation of free-living nematodes in Clade II and early branching Chromadorea^37,59,60^. Determination of the full complement of Hox genes in recently available nematode genomes across the phylum is fundamental to understanding the link between the genomic and morphological evolution of these animals. We have re-visited the Hox gene complements of Nematoda not only to understand the evolution of this key set of body pattern genes using newly available high-quality genomes but also to classify the loci into their orthologue groups as well as to look for linkage between loci in a broader phylogenetic context.

Nematodes are renowned for their simple and highly conserved body plan. However, at the sequence level, they have been shown to possess rapid homeodomain sequence and genome evolution which might be a general feature of Nematoda Hox genes^29^. We observe that it is difficult to fully prove the absence of a gene and that the missing Hox genes in some genomes could indeed be truly missing from these genomes or they could be a result of extreme homeobox sequence divergence such that the sequence similarity search methods used in this study could not detect such high sequence variability. In *C. elegans*, a reduced Hox gene complement (6 genes in 4 orthology groups) has been reported^29^ and we confirmed this reduction in this study (Figure 2), illustrating the robustness of our methods. In *Syphacia muris, Enterobius vermicularis* as well as the common ancestor of Clade IV-V nematodes, an independent loss of Hox3 had been reported^28^ and we confirmed this loss with our methods (Figure 3). This further supports the fidelity of the analysis methods employed here for detecting gene presence and absence in assembled genomes.

We have expanded the availability of Hox gene complements for a fairly large number of nematode species that were previously missing due to a lack of genomic resources. The Hox complement across the phylum Nematoda is shaped by key gene losses. We found a recent duplication of nob-1 in the model genus. This case illustrates the potential for new in-paralogues to evolve in certain lineages. In *C. elegans*, loss of nob-1 and php-3 functions leads to severe abnormalities during embryonic development, affecting both the arrangement of posterior structures and the movement of the posterior hypodermis. Additionally, this disruption results in changes from posterior to anterior cell identities and ultimately leads to non-viability^25^. Further examinations are required to establish if these roles persist across different *Caenorhabditis* species that possess nob-1.

We found presence of Hox3 in Clades I-III of Nematoda and absence in both Clade IV and V of Nematoda and Nematomorph genomes analyzed here (Figure 2). The pattern of presence and absence in Nematoda can be explained by an independent derivation in the respective Clades. Within the insect counterparts, orthologs of Hox3 (zerknüllt, zen) have undergone repeated functional alterations, resulting in the relinquishment of their conventional Hox role and subsequent reassignment of their functional domain from embryonic to extraembryonic tissues. In fact, all known zen genes have been found to participate in the development of extraembryonic membranes (EEMs). These EEMs serve to shield embryos from external environmental challenges, and their formation has enabled insects to deposit eggs in diverse habitats, ultimately facilitating their successful colonization of terrestrial environments^61^. Functional investigations are necessary to clarify the function of Hox3 in nematodes that possess this gene.

Previous analysis using transcriptome datasets and PCR surveys identified loss of ant-1 locus in the Enoplea species *Enoplus brevis*, the chromadorea *Plectus sambesi, S. muris*, the ancestor of Tylenchomorpha, the diplogasteromorph *P. pacificus*, and the ancestor of *Caenorhabditis*^28^. We now certify this pattern of presence-absence using currently available high-quality genomes from these species and further extend the loss of ant-1 locus in the pinworm *E. vermicularis* (Clade III) additionally. In *B. malayi*, it has been demonstrated that the antennapedia gene’s positioning occurs after the Bm-egl-5 Hox gene. Moreover, their homeodomain exons are alternatively joined to the same 5’ exon through cis splicing and this arrangement might signify an intermediate stage in the potential loss of Hox genes due to redundancy in other nematode species^29^. Furthermore, ant-1 was identified in *C. monodelphis* in contrast to prior PCR survey-based studies that reported its absence^28^. The presence of ant-1 exhibited variability in both Strongyles and Oscheius. Nevertheless, additional investigation is required to substantiate this distribution pattern.

We found Hox2/pb, Hox5/scr and Hox6-8/Ubx/AbdA missing in Nematoda (Figure 2). The missing genes in newly sequenced nematode genomes further validate the previous findings from PCR-based studies^28,29^. The finding of Hox2/pb and Hox6-8/AbdA in Nematomorpha (Figure 2), which are closest relatives to Nematoda, indicates that these Hox loci were lost independently in the lineage leading to Nematoda. Recent work suggests much more gene loss has occurred in metazoan genome evolution than previously thought^30,62,63^, and Hox gene loss has been observed in several other lineages^28^. The absence of Hox5/Scr in both Nematoda and Nematomorpha not only suggests an inheritance characteristic from the common ancestor of all extant nematodes but also a potential example of evolutionary conservation.

Why some Hox genes were lost early on in the evolution of the phylum and others were lost or duplicated later on remains unclear. We found that some nematodes possess Hox3 and antennapedia-like Hox gene ant-1 which belong to orthology groups present in other metazoan animals but are absent in *C. elegans* and some of its relatives (Figure 2). The loss may be due to a more specialised function that is only needed in certain nematode species or developmental contexts. For instance, Hox3 or Hox6-8/Antp may be involved in the development of specialised structures that are unique to those non-model nematode species and if these structures are lost during the course of evolution, the respective Hox genes may also become dispensable and eventually lost in some nematode lineages. In contrast, other Hox genes such as ceh-13 which is critical for embryogenesis and influences cell fate specification beyond the anterior body part^25,64^ and php-3 which regulates specification of the tail^65^ might have more essential functions that are needed for proper development of nematodes. These genes are less likely to be lost because their loss would have a more significant impact on the nematode’s development and survival. Therefore, the Hox complement in Nematoda is shaped by gene loss and might have played an important role in the evolution of their simplified body plan.

While the *C. elegans* Hox cluster is dispersed, we found intact and not dispersed Hox clusters in non model nematode genomes (Figure 4). We found rearrangement, translocation, and disruption by inversion with respect to the transcriptional orientation of the Hox genes in the clusters of analysed nematode genomes (Figure 4). This pattern of mixed arrangement reflects a highly dynamic and complex history of nematode Hox cluster evolution and organisation similar to a situation observed in *Drosophila melanogaster* where the Hox cluster is broken into parts (the Antennapedia (ANTC) and bithorax (BXC) complexes). It exhibits aberrations in the transcriptional orientation of some genes as well as both interspersed genes of independent origin and Hox-derived genes that have evolved novel developmental functions^11^. Additional Hox cluster rearrangements have also been reported in other *Drosophila* species^16,22,66,67^ as well as in the silk moth *Bombyx mori*^68^. The rearrangement of Hox genes in the Cluster of *C. elegans* has also been reported before^29^. This further suggests that the physical orientation and rearrangement of the Hox genes in the clusters are not crucial for normal nematode development and body plan conservation. The dynamics in the organisation of the nematode Hox clusters could be further evidence that strong evolutionary constraints are rather acting on the highly conserved functional domains of the nematode Hox genes to maintain the conserved body plan. Hox gene rearrangements may drive evolutionary changes causing alterations in the pattern of Hox gene expression during development. They could facilitate the evolution of new morphological structures that are characteristic of different species thereby contributing to the process of nematode speciation. As the complete genome sequences of more nematode species become available, the molecular phylogenetics of Hox cluster evolution will become clearer with an emphasis on understanding the evolution of the regulatory modules that partition Hox expression domains.

We identified non Hox genes between Hox loci (Table 1) which may act as Hox cluster coregulator candidates. Previous works identified T-box proteins as being particularly amplified in Rhabditid nematodes^69,70^. Additionally, the presence of “foreign” genes in Hox clusters of nematodes are indicators that the distances between Hox genes are quickly evolving^71^. The identification of potentially unified Hox clusters in species occupying phylogenetic positions close to or in the ancestral Clades is an indicator that perhaps the common ancestor of all nematodes had a single Hox cluster of at least six Hox genes as observed in Trichuridae (*Trichuris suis*) and *Plectus sambesii* and that these clusters have undergone remodification in different nematode lineages as a result of gene loss and Hox paralog specific duplication. The Nematoda Hox cluster organisation is highly dynamic. If present at all, the arrangement of genes in a cluster can be interrupted, shuffled, or even completely disintegrated. Further knowledge on topological Islands in the analysed nematode Hox gene clusters herein awaits an investigation.

In contrast to the nematode body plan’s conservation, essential bilaterian regulators like Hox genes have undergone significant evolutionary flexibility. The loss of some of these genes in the lineage leading to the Ur-Nematode might have played a role in roundworm body plan evolution. While the *C. elegans* Hox cluster is dispersed, we found species including *Plectus sambesii, Trichuris suis* in which the cluster is intact and not dispersed. The rearrangement of Hox genes in nematode Hox clusters may have underlying effects including the breakage of the usual colinearity of Hox gene expression in Nematoda. Further research regarding nematode Hox gene complements that we have identified in the present study should seek to understand whether and where the colinear expression is retained in the phylum. As more species become amenable to molecular methods^48,51,72^, this can be done with tools like in situ hybridization, single-cell RNA-Seq studies, and CRISPR-guided gene knockouts. In summary, our analyses suggest the potential for new and exciting work on Hox genes and regulation of Hox clusters in a broad sampling of Nematoda. New lineages with unified Hox gene clusters will likely emerge as more complete high-quality genomes become available.

## Methods

### Genome data

We included in our analysis 80 nematode genomes, including newly sequenced genomes from free-living Enoplia (Clade II) and other Chromadoria (outside Clade III-V), lineages that previously lacked genomic resources (see Figure 1). New draft genomes were generated at both the Sanger Institute and the WormL∼ab laboratory at the University of Cologne (see Supplementary Table 1). We included four genomes from Nematomorpha, *Acutogordius australiensis, Nectonema munidae, Gordionus montsenyensis*, and *Gordius aquaticus*, as the natural outgroup (see Supplementary Table 1).

### Hox Homeodomain identification

We collated a reference dataset of *C. elegans*, and *Branchiostoma floridae* (amphioxus) Hox homeodomain proteins from HomeoDB^73^. In the framework of the highly conserved Hox orthologues, the choice of these species allowed us to search with the reduced Hox gene complement of *C. elegans* as an ingroup and the full amphioxus set as an outgroup. We performed a sequence similarity search using the BITACORA^74^ gene family analysis pipeline in both genome and protein mode with default parameters (e-value of 10-5). The output of the first round of searching was used to generate an HMMER^75^ search profile containing orthologous Hox genes from multiple nematode species. Final predicted proteins and corresponding gene annotation files for each genome were generated through a second round of Hox gene orthologue search in the BITACORA environment using the HMM models. The predicted proteins were aligned with MAFFT^76^ and corresponding homeodomain sequences were visualised and extracted using Aliview^77^. Phylogenetic trees were reconstructed using the homeodomain protein sequences by maximum likelihood implemented in IQTREE2^78^. The final trees were visualised using Figtree (http://tree.bio.ed.ac.uk/software/figtree/). (See Figure 5 for visualized analysis pipeline).

**Figure 5.**
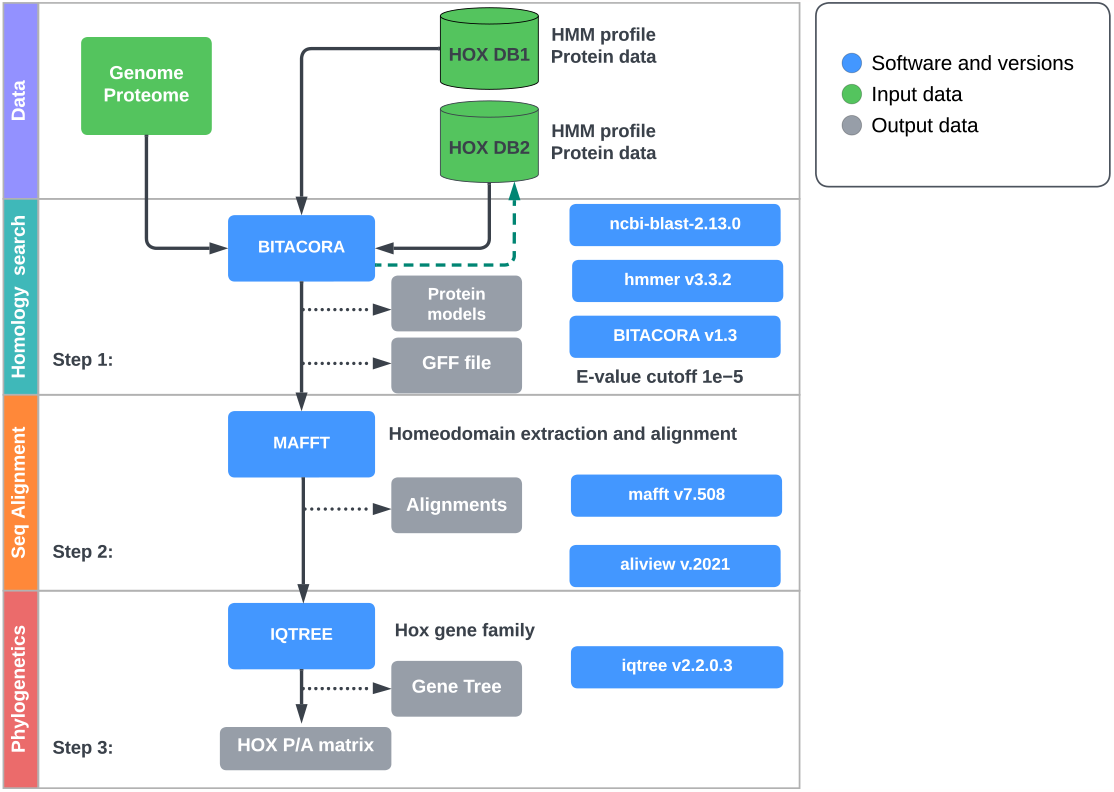
Hox gene identification workflow. The initial inputs of genome or proteome in fasta format and Hox gene reference profiles from the HOX database are colour-coded green. software tools (and versions) are colour-coded blue while output files are colour-coded grey.

### Hox cluster annotation and gene identification

After identification of HOX homeodomains, we extracted the respective scaffolds or contigs for annotation of complete Hox genes in recently available genomes without genome annotation. To search for potential clusters of Hox genes in assembled genomes, genomic scaffolds or contigs containing Hox genes were extracted from the respective assemblies using seqtk (https://github.com/lh3/seqtk). We used AUGUSTUS (v3.5.0)^79^ to predict complete protein models in contigs and scaffolds. To improve the accuracy of gene prediction AUGUSTUS was trained on *C. elegans* and a reference set of Hox gene profiles from *C. elegans* and *B. floridae* as hints for Hox gene finding. A database of predicted proteins from AUGUSTUS was constructed for search using BLASTP (v2.9.0)^80^ with an e-value of 10-5 using a query file containing a reference set of Hox protein sequences. Hox gene loci were manually inspected if they were in clusters (We define a Hox cluster as a group of four or more Hox genes spanning a genomic loci with a size less than the size of the loosely connected *C .elegans* Hox genes (i.e less than 0.3 kb) on the same contig or scaffold. To validate the presence of Hox clusters we used HbxFinder pipeline available at GitHub (https://github.com/PeterMulhair/HbxFinder) to identify homeobox genes in the respective contigs or scaffolds. Hox gene cluster organisation and strand orientation maps from the AUGUSTUS output GFF files were drawn using Affinity Designer 2 (https://affinity.serif.com/en-gb/learn/designer/desktop/). Presence of non Hox genes were extracted and counted from the respective GFF structural annotation files.

## Acknowledgements

This work was funded by a DFG ENP grant to PS (grant number: 434028868). Work at the Wellcome Sanger Institute was funded by Wellcome Trust Grants 206194 and 218328. For the purpose of Open Access, the author has applied a CC BY public copyright licence to any Author Accepted Manuscript version arising from this submission.

## Author contributions statement

K.J, S.P and B.M. conceptualized revisiting Hox genes in recently available genomes across the Nematoda, A.A. and B.A. conducted the experiment(s), K.J. and L.D. analysed the results. S.P and H.A supervised the analysis. K.J, S.P, K.A and L.S prepared and provided the draft genomes. S.P, B.M and H.A checked through the manuscript and provided critical proofreading of the manuscript. All authors reviewed the manuscript.

## Supplementary Information

Sources of data used in this article are listed in Table 1 (Supplementary information). The already published genomes are publicly available but the draft genomes can be accessed on request. Custom scripts, command lines, and all results from these analyses are available at github (https://github.com/jkirangw/NematodeHOX.git).

